# Genes identified in rodent studies of alcohol intake are enriched for heritability of human substance use

**DOI:** 10.1101/2021.03.22.436527

**Authors:** Spencer B. Huggett, Emma C. Johnson, Alexander S. Hatoum, Dongbing Lai, Jason A. Bubier, Elissa J. Chesler, Arpana Agrawal, Abraham A. Palmer, Howard J Edenberg, Rohan H.C. Palmer

**Author notes:** Corresponding Author: Spencer B. Huggett, Behavioral Genetics of Addiction Laboratory – Department of Psychology, Emory University, 36 Eagle Row, Atlanta, GA 30322, USA.

## Abstract

**Background:** Rodent paradigms and human genome-wide association studies (GWASs) on drug use have the potential to provide biological insight into the pathophysiology of addiction.

**Methods:** Using GeneWeaver, we created rodent alcohol and nicotine gene-sets derived from 19 gene expression studies on alcohol and nicotine outcomes. We partitioned the SNP-heritability of these gene-sets using four large human GWASs: 1) alcoholic drinks per week, 2) problematic alcohol use, 3) cigarettes per day and 4) smoking cessation. We benchmarked our findings with curated human alcoholism and nicotine addiction gene-sets and performed specificity analyses using other rodent gene-sets (e.g., locomotor behavior) and other human GWASs (e.g., height).

**Results:** The rodent alcohol gene-set was enriched for heritability of drinks per week, cigarettes per day, and smoking cessation, but not problematic alcohol use. However, the rodent nicotine gene-set was not significantly associated with any of these traits. Both rodent gene-sets showed enrichment for several non-substance use GWASs, and the extent of this relationship tended to increase as a function of trait heritability. In general, larger gene-sets demonstrated more significant enrichment. Finally, when evaluating human traits with similar heritabilities, both rodent gene-sets showed greater enrichment for substance use traits.

**Conclusion:** Our results suggest that rodent gene expression studies can help to identify genes that capture heritability of substance use traits in humans, yet the specificity to human substance use was less than expected due to various factors such as the genetic architecture of a trait. We outline various limitations, interpretations and considerations for future research.

## INTRODUCTION

Alcohol consumption, alcohol use disorder (AUD), cigarette smoking and smoking cessation are all complex, heritable phenotypes, with twin and family estimates of heritability ranging from 20-70%^1,2^ Molecular genetic studies have demonstrated that all of these behaviors are highly polygenic^3^, meaning that many common genetic variants of small effect sizes contribute to the variation in these phenotypes. Recent genome-wide association studies (GWASs) of these behaviors have leveraged collaborative efforts and increasing sample sizes (currently ranging from ∼40,000 to over a million individuals) to identify multiple genetic loci^4–7^.

To prioritize the potentially relevant genes of the identified risk loci for experimental follow-up, researchers often integrate functional data (i.e., transcriptomic, epigenetic, and chromatin interaction data) or test whether associated genes are enriched among curated gene-sets like the Kyoto Encyclopedia of Genes and Genomes (KEGG^8^). KEGG gene-sets and pathways are created via computational and manual methods, primarily derived from experimental evidence in model organisms^9^. They have provided a valuable resource to enhance the biological understanding of complex human traits^10^ and inform therapeutic targets for alcohol use disorder^11^, but these pathways are thought to be incomplete and lack both tissue specificity and behavioral nuance^12^.

Animal paradigms have helped characterize the underlying neurobiological components of addiction and its basic behavioral processes, but the extent to which they identify the same genes that influence the genetic propensity for human substance use and use disorders remains unclear. Neurobiological facets of addictive behaviors are likely to be shared across mammalian species and drugs of abuse; thus, integrating evidence from rodent genetic studies with human GWAS could help prioritize human GWAS signals that may represent conserved aspects of the addiction pathway^13^. Unique challenges exist for cross-species data integration, including homology of genes and the substantial differences between human and model organism phenotypes. There is an urgent need to understand whether and under what conditions genes identified in model organism genomic studies of addiction are also implicated in human GWASs of related traits^14^. Few studies have systematically examined the overlap of genes identified in rodent paradigms that model aspects of addiction^15–18^ with those identified in humans. Prior studies along these lines have focused on individual genes^19^; polygenic cross-species approaches are limited (although one recent study used genome-wide complex trait analysis and polygenic score approaches and found evidence of cross-species genetic overlap for nicotine consumption^20^).

The goals of our study were to: 1) investigate the contribution of homologous rodent alcohol and nicotine gene-sets to the genome-wide SNP heritability (*h*^2^_SNP_) of human alcohol and tobacco use and related phenotypes, 2) assess the specificity and sensitivity of these results using KEGG pathways, non-substance related gene-sets, and non-substance related GWASs and 3) explore the individual behavioral paradigms that best capture genetic variation in human substance use-related traits. To do this, we used GeneWeaver^21^, a cross-species functional genomics database, to identify gene-sets from brain-related gene expression studies of alcohol and nicotine consumption, exposure, and selective breeding paradigms in various mouse and rat populations. We hypothesized that rodent substance use gene-sets would capture relevant and specific genetic variation for human alcohol and tobacco use-related traits. For an overview of our study, see **Figure 1**.

**Figure 1.**
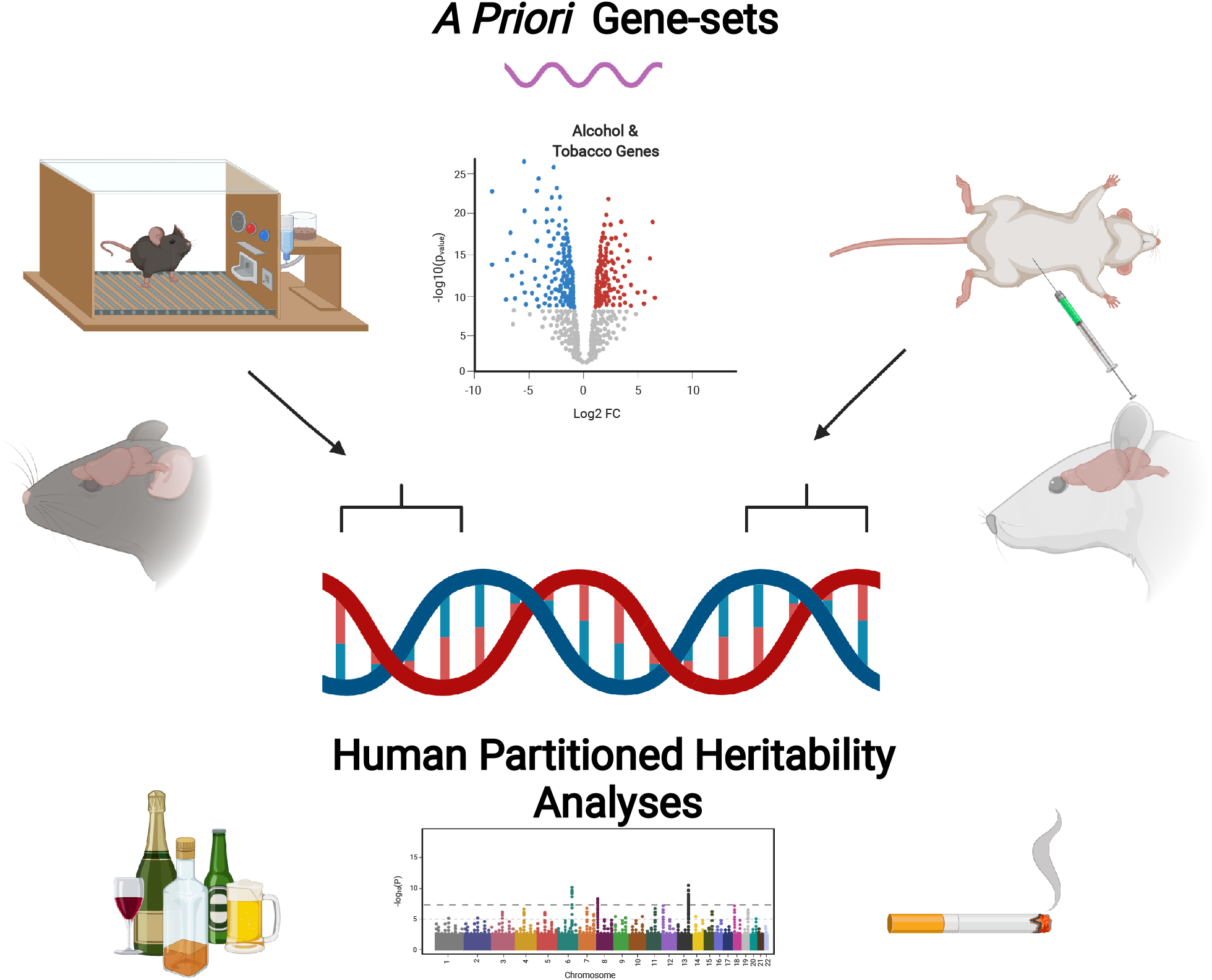
Schematic of our study. Image created via Biorender.com

## MATERIALS AND METHODS

### A Priori Gene-Sets

We queried GeneWeaver’s^21^ database of heterogeneous functional genomics data (https://www.geneweaver.org/) to identify six gene-sets. Specifically, we used curated publication-derived gene-sets (Tier 3; queried July 2020; search terms: ethanol, alcohol, nicotine and tobacco) that included differential gene expression from mouse and rat brain regions following alcohol or nicotine intake, exposure, and selective breeding experiments, as well as two negative control studies that are described below. The homologous human genes corresponding to the rodent genes were identified using biomaRt^22^; we only used genes on autosomal chromosomes because the human substance use GWASs did not include results from the sex chromosomes. In total, we used gene-sets derived from 21 studies, involving a total of 20 inbred or outbred genetic backgrounds of mice and rats (n=750, see **Table 1**; and **Supplementary Table S1** for more details). Of these 21 studies there were 17 that focused on alcohol, identifying 4,310 genes in total. To prioritize the most reliably associated genes, our analyses focused on the 828 genes observed in at least two of these studies. We found two studies that assessed rodent nicotine outcomes that identified 417 genes. Due to minimal overlap of genes across the two nicotine studies, all genes were used. The last two studies foucused on non-substance gene-sets that are popular reference paradigms to isolate specific aspects of drug use from the animal literature: sucrose consumption (87 genes^23^) and locomotor behavior (546 genes^24^).

**Table 1.** Gene-sets for Alcohol, Nicotine and Non-Substance Use Behavioral Paradigms.

| **Alcohol Gene-set** |  |  |  |  |  |
| --- | --- | --- | --- | --- | --- |
| GeneWeaver_ID | PMID | Species | Behavior | Brain_Regions | # Genes |
| GS127422; GS127424; GS127426 | 18397380 | Mouse (B6J vs B6C mice) | Naïve Alc Consum/Pref | Cortex; Hipp; Cerebell; Str | 44 |
| GS14962-65 | 15597075 | Mouse (B6J vs DBA/2J mice) | Naïve Alc Consum/Pref | Amy; Str; Hipp | 69 |
| GS216937 | 23550792 | Mouse (HDID vs HS/NPT mice) | Naïve Binge Consum | vStr | 68 |
| GS14921-GS14924 | 15002731 | Mouse (B6J / DBA/2J mice) | Acute Exposure | Whole brain | 74 |
| GS127341-GS127346 | 21223303 | Mouse (B6J mice) | Binge Drinking | Str; PFC; VTA; Cerebell; Hipp | 1393 |
| GS246643; GS246644 | 26482798 | Mouse (B6J x FVB/NJ F1 mice) | Binge Drinking | VTA | 1086 |
| GS14925; GS14926 | 12886948 | Mouse (B6J / DBA/2J mice) | Alc Withdrawl | Hipp | 32 |
| GS1777 | 12805289 | Mouse (HAFT vs LAFT mice) | Acute Tolerance | Whole brain | 129 |
| GS14945-GS14948 | 17517326 | Rat (iP vs iNP rats) | Naïve Alc Consum/Pref | NAC; Amyg; PFC; Str; Hipp | 64 |
| GS246645; GS246646 | 26455281 | Rat (Warsaw, Low vs High Pref) | Naïve Alc Consum/Pref | NAC; Hipp; PFC | 215 |
| GS75752 | 15846778 | Rat (AA, P, ANA Rats) | Naïve Alc Consum/Pref | PFC | 42 |
| GS74017-GS74020 | 15052257 | Rat (AA, ANA Wistar Rats) | Naïve Alc Consum/Pref | ACC; NAC; Amyg; Hipp | 44 |
| GS75753 | 11772933 | Rat (Wistar) | Chronic Intermittent Alc | ACC & Amyg | 28 |
| GS18838; GS18840 | 12462420 | Rat (Lewis Rats) | Chronic Exposure | Hipp | 65 |
| GS37147 | 19666046 | Rat (P rats) | Binge Drinking | Nac | 345 |
| GS14929; GS14930 | 18405950 | Rat (iP Rats) | Binge Drinking | NAC; Amyg | 266 |
| GS246761 | 27061086 | Rat (P rats) | Binge Drinking | Periaqueductal gray | 1411 |
| **Rodent Nicotine Gene-set** |  |  |  |  |  |
| GeneWeaver_ID | PMID | Species | Behavior | Brain_Regions | # Genes |
| GS14888-GS14893 | 17504244 | Mouse (B6J & C3H/HeJ Mice) | Chronic Exposure | PFC; Nac; Hipp; Amyg; VTA | 374 |
| GS14885 | 17234382 | Rat (Long Evans Rats) | Chronic Exposure | PFC; Str; Hipp | 65 |
| **Non-substance Gene-sets** |  |  |  |  |  |
| GeneWeaver_ID | PMID | Species | Behavior | Brain_Regions | # Genes |
| GS36166-GS36193 | 19958391 | Mouse (BXD Mice) | Locomotor Behavior | Str; Cortex; Cerebell; Hipp; Whole Brain | 546 |
| GS14885 | 8405950 | Rat (iP Rats) | Sucrose Consumption | Nac; Amyg | 87 |

To benchmark our findings, we also investigated two curated Kyoto Encyclopedia of Genes and Genomes (KEGG) Pathways^8^. Specifically, we examined the “Human Nicotine Addiction” (total genes = 36) and “Alcoholism” (total genes = 95) pathways, to see if they accounted for a significant amount of variance in the genetic predisposition of alcohol and tobacco use traits. For a summary of all genes used in our partitioned heritability analyses (and their overlap), see **Figure 2** and **Supplementary Table S2**.

**Figure 2.**
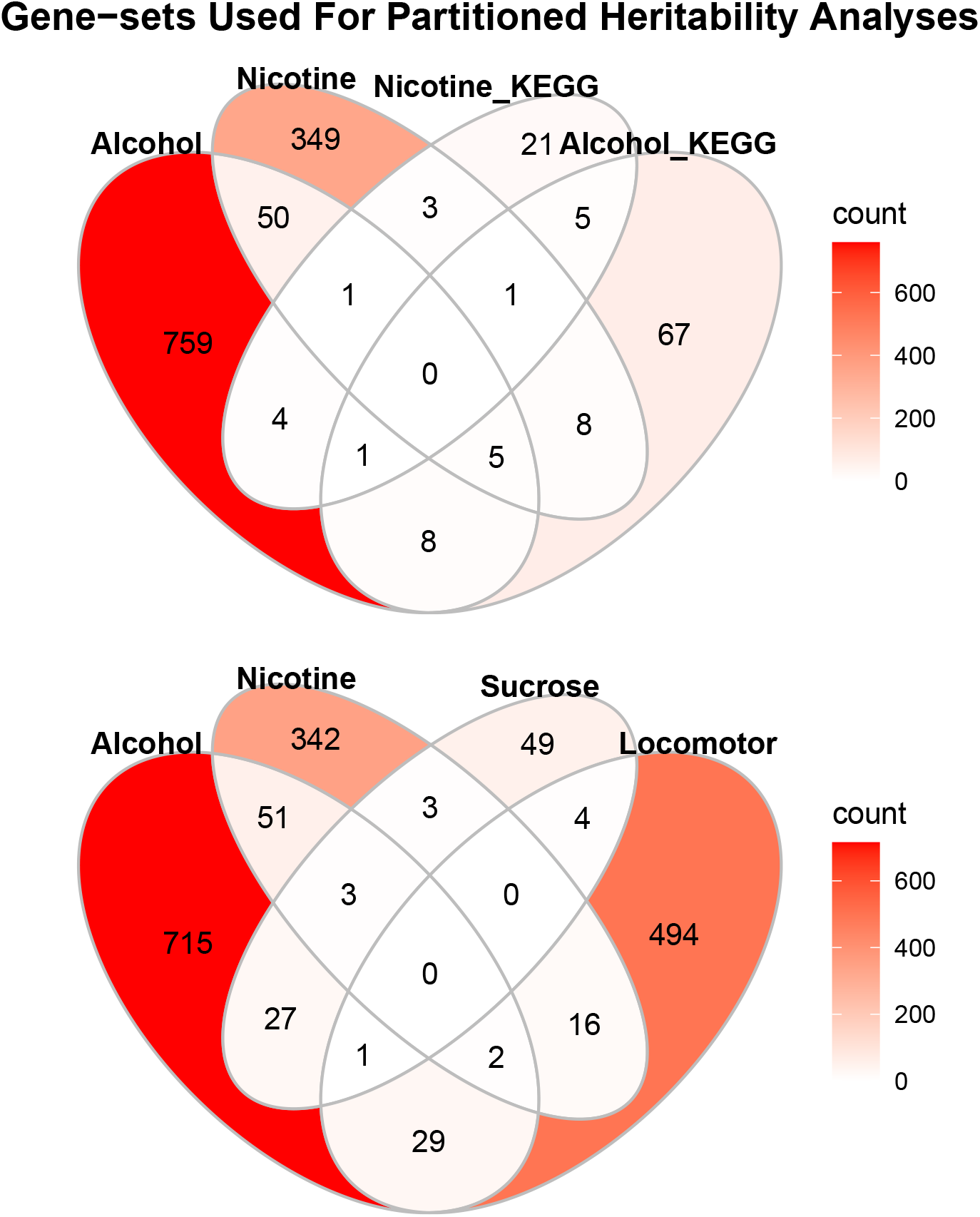
A Priori Gene-sets Used in Partitioned Heritability Enrichment Analyses. Alcohol, Nicotine Sucrose and Locomotor gene-sets were derived from GeneWeaver and Alcohol_KEGG and Nicotine_KEGG gene-sets represented the Human Alcoholism and Nicotine Addiction genes, respectively. Note these plots only include genes that were homologous with humans and located on human autosomal chromosomes. Alcohol rodent genes were required to replicate across at least one other study.

### Human Substance Use GWAS Summary Statistics

The relevance of the identified gene-sets were examined using summary statistics from several GWASs of European ancestry including samples from the United Kingdom BioBank, Million Veteran Program and the GWAS and Sequencing Consortium of Alcohol and Nicotine Use (GSCAN). These are among the largest studies to date for each of the alcohol and nicotine related measures. Note that these samples did not include the 23andMe data, since these data are not publicly available. The individual GWASs included demographic covariates (age, sex, 5-10 ancestral principal components). The data source and sample size for each measure are as follows:

#### Problematic alcohol use (PAU)

Summary statistics for problematic alcohol use were derived from a meta-analysis by Zhou et al^25^ (N =435,563) collapsing across alcohol dependence^26^, alcohol use disorder^27^, and Alcohol Use Disorder Identification Test-Problem items^28^.

#### Drinks per week (DPW)

Drinks per week summary statistics came from a GWAS on self-reported drinks per week^5^ (N = 537,349).

#### Cigarettes per day (CPD)

We used summary statistics from a GWAS on smokers’ self-reported average number of cigarettes smoked per day^5^ (N = 263,954).

#### Smoking cessation (Cig_Cessation)

Summary statistics for current vs. former smoking status were derived from the GSCAN GWAS^5^ (N = 312,821).

### Partitioned Heritability Analyses

We performed partitioned heritability analyses for each gene-set using Linkage Disequilibrium Score Regression (LDSC)^29,30^. Partitioned LDSC produces an estimate of whether single nucleotide polymorphisms (SNPs) within and around the genes in a given gene-set account for a significant proportion of the heritability of a given trait relative to the number of variants included. Specifically, enrichment is calculated as: (proportion of SNP-heritability explained)/(proportion of SNPs in gene-set relative to all other SNPs). We tested for heritability enrichment using the six gene-sets described in the previous sections: 1) rodent alcohol (Alcohol) 2) rodent nicotine (Nicotine), 3) alcoholism (Alcohol_KEGG), 4) human nicotine addiction (Nicotine_KEGG), and two non-substance use traits: 5) rodent locomotor behavior (Locomotor) and 6) rodent sugar consumption (Sucrose). We used LDSC’s default gene window size (100 kb) to assign SNPs to genes based on the expectation that up to 80% of local gene regulatory regions occur within 100 kb of a gene^31^. Given recent evidence suggesting a decrease in enrichment beyond 10 kb for CPD^20^, we also report results from a smaller 10 kb window. To determine significance, our study adjusted p-values with a Benjamini-Hochberg False Discovery Rate (BH-FDR; FDR < 5%).^32^

### Sensitivity analyses

To examine the specificity of the GeneWeaver gene-sets and to provide context for interpretation, we also tested the heritability enrichment of the gene-sets in 14 non-substance use GWASs with a range of heritabilities and genetic correlations with substance use traits (see **Table 2** and **Supplementary Information**). These included neurological and neuropsychiatric traits (e.g., schizophrenia, Alzheimer’s disease) as well as anthropometric, cardiometabolic, and other traits that are theoretically unrelated to drug use (e.g., height, wearing glasses).

**Table 2.** Human GWASs Used in Partitioned Heritability Analyses.

| Substance Use GWASs |  |  |  |
| --- | --- | --- | --- |
| GWAS | PMID | N | $h^2_{\text{SNP}}$ |
| Drinks per Week | 18397380 | 537,349 | 4 |
| Problematic Alcohol Use | 15597075 | 435,563 | 7 |
| Cigarettes per Day | 23550792 | 263,954 | 8 |
| Smoking Cessation | 15002731 | 312,821 | 5 |
| Non-Substance Use GWASs |  |  |  |
| GWAS | PMID | N | $h^2_{\text{SNP}}$ |
| Side of Head Phone Used | NA | 303,009 | 5 |
| Tinnitus | NA | 117,882 | 12 |
| Wear Glasses | NA | 360,677 | 4 |
| Ankle Width | NA | 206,589 | 30 |
| Hip Circumference | NA | 360,521 | 22 |
| Height | NA | 360,388 | 49 |
| Alzheimers | 30617256 | 455,258 | 33 |
| ASD | 28540026 | 173,773 | 34 |
| ADHD | 30478444 | 55,374 | 26 |
| MDD | 29700475 | 480,359 | 9 |
| Schizophrenia | 25056061 | 79,845 | 46 |
| Bone Mineral Density | NA | 206,496 | 32 |
| Heart Disease | NA | 361,194 | 16 |
| Diabetes | NA | 360,192 | 21 |

### Exploratory analyses

We performed secondary analyses to evaluate patterns within our data. Because the majority of the rodent studies we used pertained to alcohol, we performed exploratory analyses to determine which alcohol behavioral paradigms corresponded with individual human substance use traits. These analyses examined mice and rat data separately and considered various categories of alcohol behavioral paradigms using a total of seven gene-sets. For mice, we examined 1) binge drinking (2,317 genes), 2) naïve strain differences (e.g., differences among strains selected for -or known to differ in -alcohol consumption and preference; 140 genes), 3) acute exposure (30 genes) and 4) tolerance + withdrawal (159 genes). For rats, we examined 1) binge drinking (1,912 genes), 2) naïve strain differences (359 genes) and 3) chronic exposure (91 genes; see **Supplementary Table S1-S2** for more information). Of the 828 genes from our alcohol analyses, 71.86% came from mouse paradigms of binge drinking. The seven categories of rodent alcohol paradigms captured largely non-overlapping sets of genes (see **Supplementary Figures S1 – S3**). Because each gene-set was derived from a smaller number of studies, we included genes that occurred in only one of the datasets, unlike our main alcohol analysis, which required rodent alcohol genes to appear in two or more datasets.

## RESULTS

SNPs in and around genes in the rodent alcohol gene-set were enriched for human drinks per week, cigarettes per day and smoking cessation (all OR > 1.39; all p < 0.003, all p_adj_ < 0.017; 10.63%-12.54% of *h*^2^_SNP_) -but not problematic alcohol use (**Figure 3**). In contrast, the nicotine gene-set did not show significant enrichment for any human substance use trait. After multiple testing correction, we found that the heritabilities of human alcohol or tobacco use were not enriched for KEGG addiction pathways, rodent locomotor behavior or sucrose consumption gene-sets (see **Figure 3** and **Supplementary Tables S3-S4**). Thus, of the six gene-sets we examined, only the rodent alcohol gene-set showed significant enrichment.

**Figure 3.**
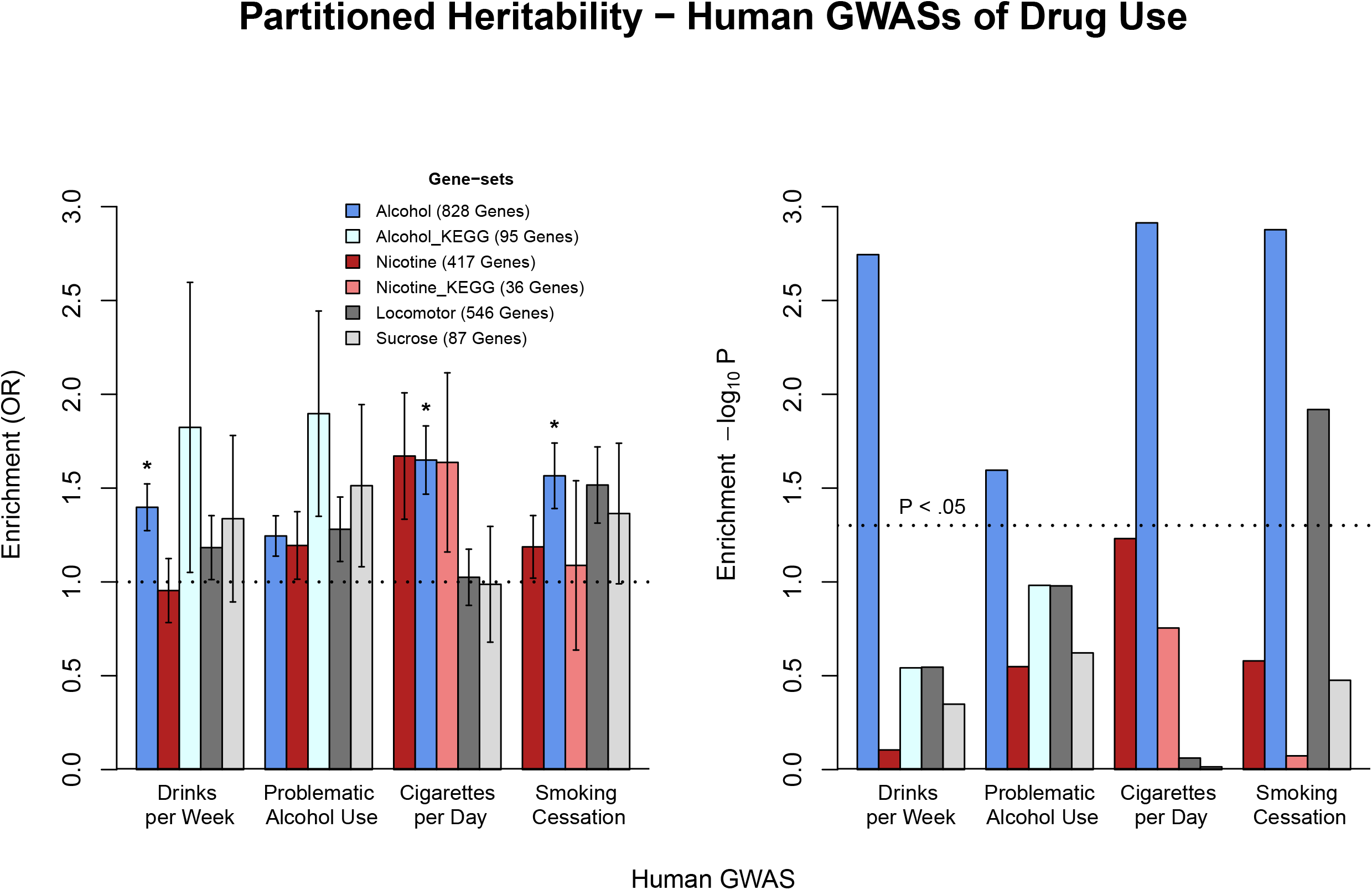
Partitioned heritability enrichment results for human substance use GWAS. The odds ratio and standard error are shown in the left panel and the –log_10_ pvalues on the right. Legend shows the gene sets and the number of genes in each geneset. * represents a significant result after using a BH-FDR correction for multiple testing (p_adj_ < 0.05).

Genetic correlations among all GWAS traits were estimated and reported in **Figure 4**. We observed significant genetic correlations between substance use GWASs and psychiatric phenotypes as well as genetic associations among tobacco traits, cardiometabolic and body morphology traits (all p_adj_ < .05). Apart from those instances, non-substance use GWASs were generally uncorrelated with alcohol and tobacco use GWASs.

**Figure 4.**
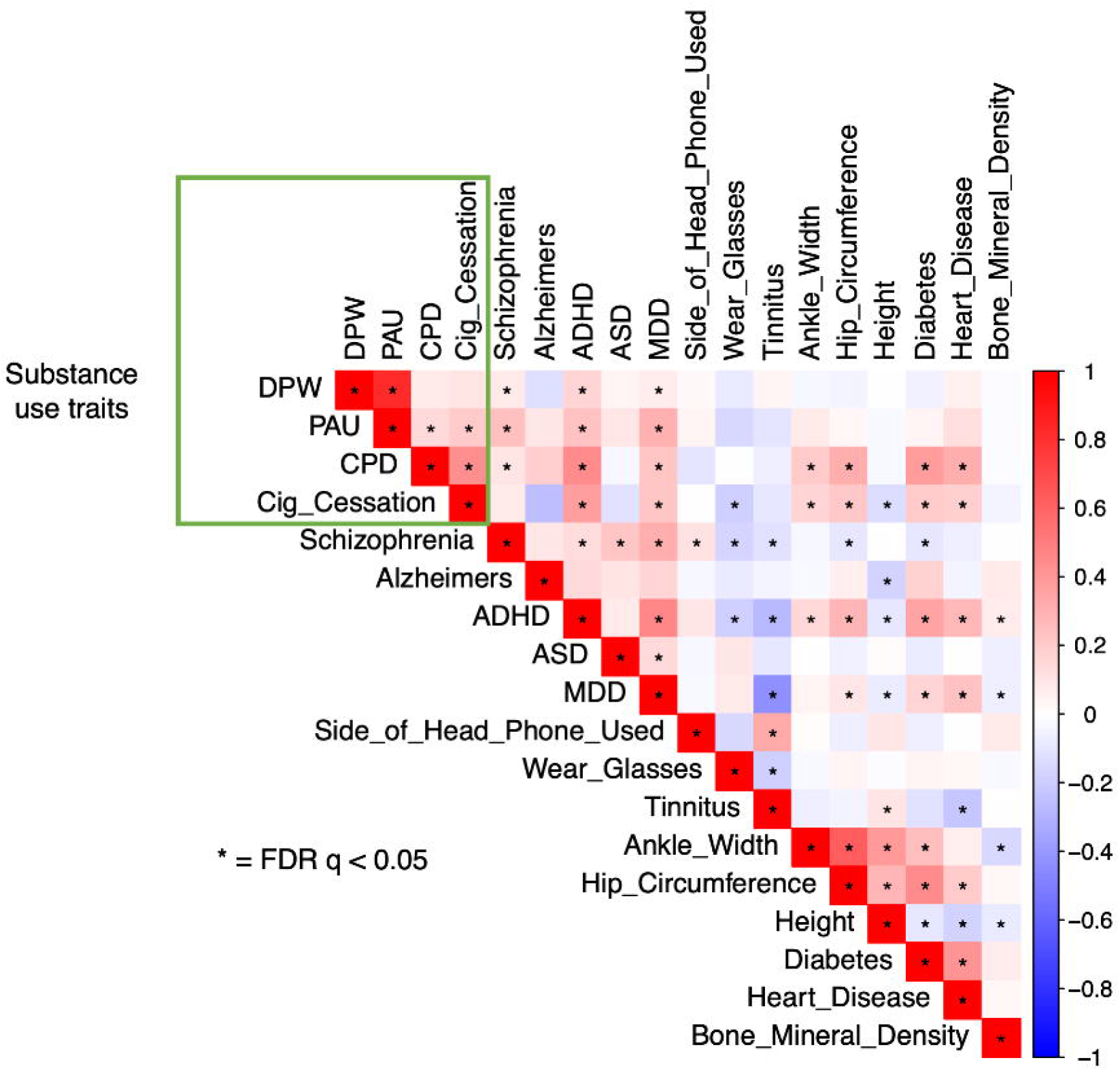
Genetic Correlations of Human GWAS Traits used in Partitioned Heritability Analyses

We tested heritability enrichment for 14 non-substance use GWAS traits with the rodent gene-sets. The heritability of height and hip circumference were enriched for genetic variants in and around genes in the rodent alcohol and nicotine gene-sets (see **Supplementary Table S5**). In addition, the rodent alcohol gene-set (but not the nicotine gene-set) contributed significantly to the heritability of schizophrenia, Alzheimer’s disease, and ankle width (see **Supplementary Table S5**). Note that, except for Alzheimer’s disease, the heritabilities of height, hip circumference, schizophrenia and ankle width were significantly enriched for LDSC’s conserved mammalian gene-sets annotation (all p < 0.006; see **Supplementary Figure S4**). Additionally, excluding hip circumference, the heritabilities of the 14 non-substance use traits were not enriched for rodent sucrose consumption or locomotor behavior gene-sets (see **Supplementary Table S6**).

When we further examined the puzzling association between rodent drug use gene-sets and non-substance use traits, we found a linear relationship between a trait’s heritability and the significance in partitioned heritability analyses (see **Supplementary Figure S5**) as well as the extent of enrichment (see **Figure 5**). These linear relationships persisted among all human GWAS traits and rodent gene-sets of sucrose consumption and locomotor behavior (see **Supplementary Figures S6 – S7**). Notably, rodent nicotine and alcohol gene-sets showed greater enrichment and more significant p-values for the heritability of human substance use than non-substance use traits of similar heritabilities (for traits with *h*^2^_SNP_ < 15%: *M*_OddsRatio_ = 1.45 vs. *M*_OddsRatio_ = 1.12, t = 2.30, p = 0.048; *M*_-log10(P)_ = 1.58 vs. *M*_-log10(P)_ = 0.49; t = 2.36, p = 0.037).

**Figure 5.**
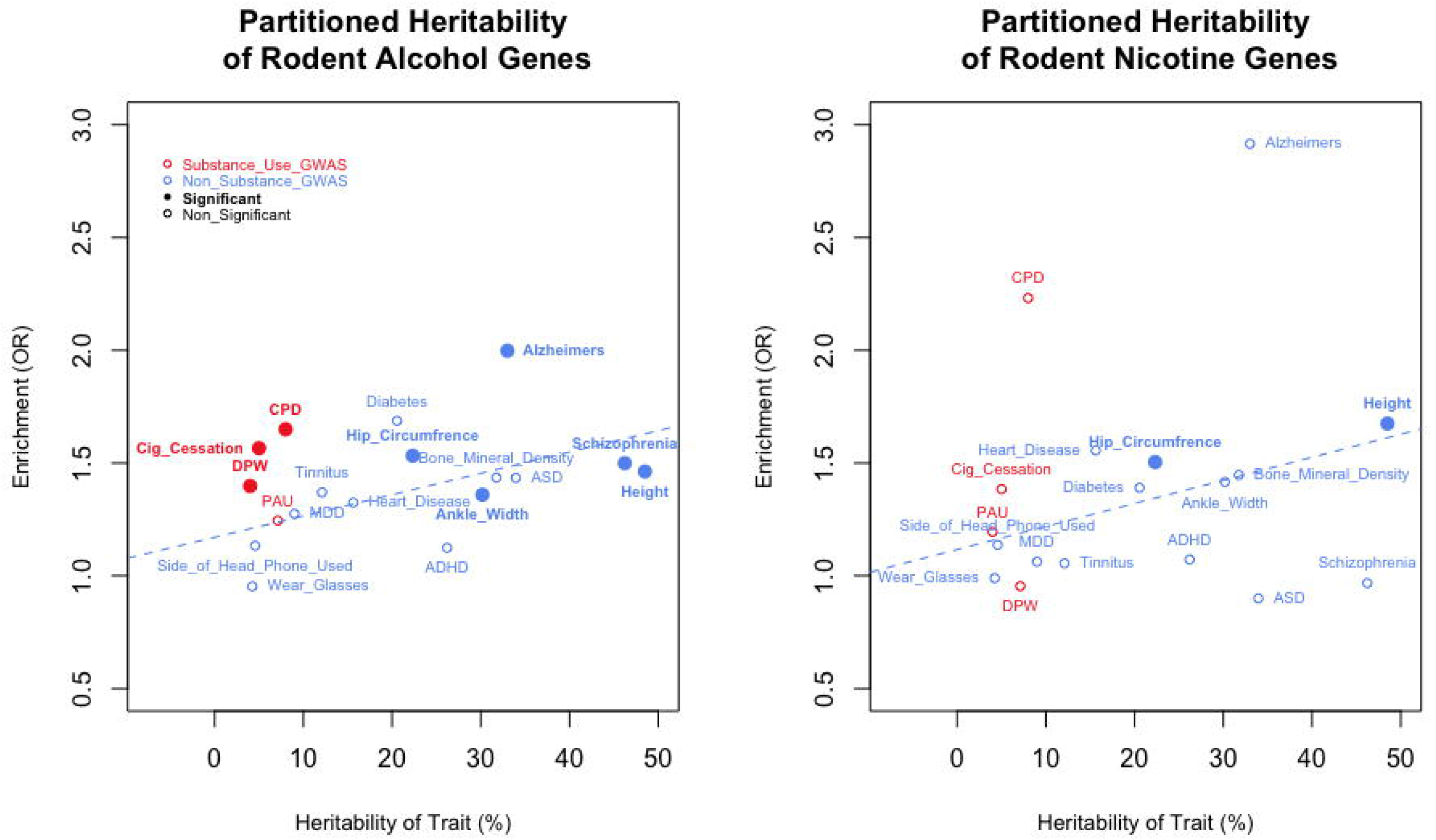
Enrichment estimates increase with heritability of trait studied. Overall, substance use traits tend to show more enrichment when compared with traits of similar heritability. The dashed blue line shows the best fitting regression line of a non-substance use trait’s heritabilty predicting the enrichment (Odd’s Ratio; OR) from partitioned heritability analyses. DPW = drinks per week, PAU = problematic alcohol use, CPD = cigarettes per day and Cig_Cessation = smoking cessation, ADHD = attention deficit hyperactivity disorder; MDD = major depressive disorder; ASD = autism spectrum disorder.

We compared the results of our partitioned heritability analyses when using the default 100 kb window with findings from a 10 kb window. Collapsing across analyses, we found that the enrichment (odds ratios; OR) and p-values did not significantly differ across 10 kb vs. 100 kb windows (all t < 1.53, all p > 0.126). We report results from our 100 kb analyses in text and also include the 10 kb findings in the supplement.

Lastly, we performed exploratory analyses to examine which of the rodent alcohol datasets were driving the enrichment of the human GWAS of drinks per week, cigarettes per day and smoking cessation. While most traits did not survive correction for multiple testing (FDR < 5%), mouse paradigms of binge drinking (e.g., the drinking in the dark model) accounted for significant genetic variance in human cigarettes per day and smoking cessation (all *p* < 0.001; all p_adj_ < 0.012; see **Figure 6**).

**Figure 6.**
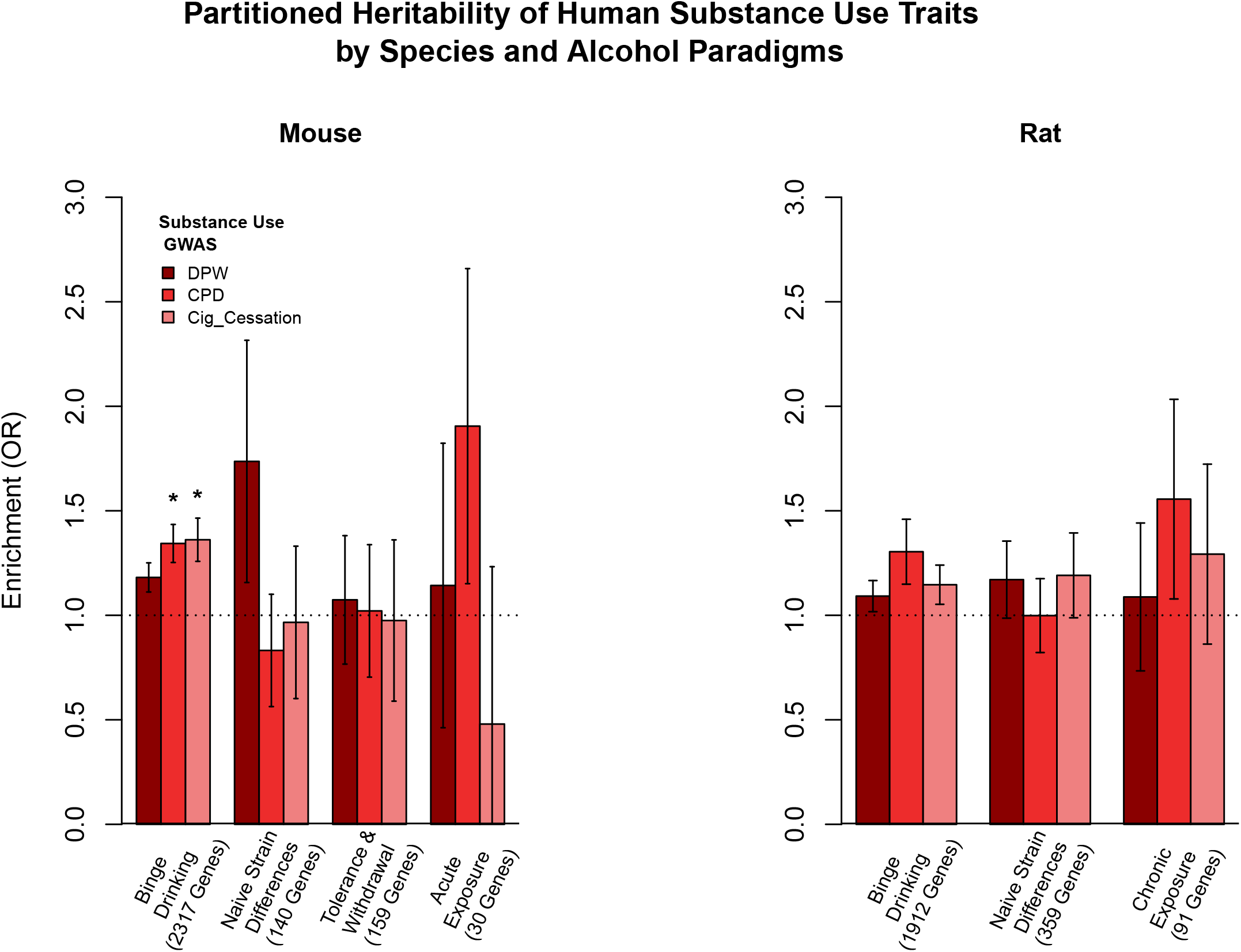
Heritability enrichment analysis for human tobacco and alcohol use, partitioned by rodent alcohol behavioral paradigm. Barplot showing the odds ratio and (standard error) from partitioned heritability analyses. Rodent behaviors were collapsed into a few categories that recapitulate different aspects of alcohol/substance use. Only mouse models of binge drinking survived correction for multiple testing as indicated by the asterisk. DPW = drinks per week; CPD = cigarettes per day; Cig_Cessation = smoking cessation.

## DISCUSSION

Our study evaluated the hypothesis that gene-sets identified in rodent alcohol and nicotine gene expression studies would be enriched for SNP-heritability in human alcohol and tobacco use traits. We found that rodent paradigms related to alcohol consumption were significantly enriched for heritability of alcohol consumption, tobacco smoking and smoking cessation, but not problematic alcohol use. These results suggest that rodent alcohol use paradigms may relate more closely to general human drug consumption than to problems arising from excessive alcohol consumption and further reinforces previously reported genetic differences between problematic alcohol use and consumption measures^26,28,33^. However, results from rodent nicotine exposure paradigms did not demonstrate enrichment in any human substance use trait. These findings are in contrast to previous cross-species partitioned heritability research that found enrichment of model organism nicotine genes in the SNP-heritbality of human cigarettes per day^20^. The current study derived a smaller nicotine gene-set from less animal paradigms, fewer species, used different sized windows surrounding genes and tested for enrichment using an alternative methodology. Notably, our rodent alcohol gene-set included a rich array of studies investigating many aspects of the biobehavioral processes of substance use including: binge consumption, non-voluntary exposure, tolerance, withdrawal and drug preference, whereas the rodent nicotine gene-set in the current study was limited to non-voluntary (experimenter-administered) drug exposure.

Specificity analyses of non-substance use GWAS revealed that genes from rodent paradigms of nicotine and alcohol use explained significant amounts of variance for seemingly unrelated human traits (e.g., hip circumference and height). There are several possible explanations for these unexpected findings. Highly heritable and highly polygenic traits were more likely to yield significant enrichment – especially for larger gene-sets. For instance, two-thirds of non-substance use traits with a h^2^_snp_ above 20% were significantly enriched for the rodent alcohol gene-set (828 genes). Our analyses revealed positive linear relationships between a trait’s SNP-heritability and partitioned heritability enrichment for all rodent gene-sets (see **Figure 3** and **Supplementary Figures S4 – S6**). Relative to the non-substance use traits with similar heritabilities (e.g., wearing glasses vs alcohol and tobacco use traits), the rodent alcohol and nicotine gene-sets demonstrated greater specificity for human alcohol and tobacco traits. It should be noted that the rodent gene-sets we used included only a subset of rodent genes with human homologs, and may therefore be identifying the subset of human genes that are better conserved across species. In support of this idea, we observed that the human traits that demonstrated enrichment for rodent drug use gene-sets were also enriched for conserved mammalian genes. Another point to consider is that the rodent drug use gene-sets were derived from brain tissues and captured relatively large genes – particularly for alcohol (alcohol: *M*_gene_size_ = 82.4 kb; nicotine: *M*_gene_size_ = 69.6 kb; all homologous genes: *M*_gene_size_ = 67.1 kb). Thus, our rodent gene-sets may be comprised of genes associated with general brain functionality and may increase power for partitioned heritability analyses by including more SNPs in and around protein-coding genes. Likewise, larger sets of genes showed a tendency to be significantly linked with a GWAS trait. Overall, our results indicate that the rodent drug use genes are not necessarily *specific* to the heritability of human substance use traits. Note that the amount of variance explained by any one gene-set was modest and consistent with polygenic architectures for complex human traits – including substance use^3,20^.

We found preliminary evidence that mouse binge drinking paradigms (drinking in the dark^34^) captured molecular mechanisms relevant for the genetic predisposition of human substance use. Binge-like consumption is a critical component of substance use escalation. However, we caution readers that these findings could be due to the large number of genes in the mouse binge drinking gene-set (2,317 genes). Combining across many behavioral paradigms, species and genetic strains generally increased prediction to a corresponding human trait (as seen in the lack of enrichment for the nicotine gene-set, which only drew from studies of non-voluntary nicotine exposure). Ultimately, the inclusion of a greater breadth and sophistication of genetic studies of rodent behavioral paradigms may provide specificity and context to mechanisms of GWAS variant action and their roles in human substance use.

There are several limitations of the current study. First, we limited our analyses to human GWASs of European-ancestry samples to maximize sample size and statistical power. Likewise, most rodent experiments rely on a small fraction of extant rodent genetic diversity due to the use of domesticated populations that have been selected for success under laboratory conditions and in some cases have been inbred strains. These constraints may limit the generalizability of our results. Second, our animal data for nicotine intake were limited to two non-voluntary nicotine exposure paradigms. Non-voluntary exposure paradigms model the physiological components of drug use, but do not explicitly model human drug use *behaviors*.

Future cross-species genetic studies may benefit from integrating multiple behavioral paradigms, strains, species, tissue types and potentially binge-like or escalated use paradigms. Nevertheless, we contend that more genes are not necessarily better; larger rodent gene-sets may contribute to false positives and a lack of specificity, especially if the heritability of a trait is already enriched for conserved mammalian genes. Finally, our study used rodent RNA findings to inform analyses about human DNA associations. Using similar data types for cross-species genetics research (e.g., rodent GWAS with human GWAS) may also demonstrate utility and increased precision.

The current analyses provide evidence that gene-sets derived from basic research in model organisms show some correspondence with human GWASs of substance use, but to a lesser extent and with less trait specificity than we initially hypothesized. As human GWAS sample sizes continue to grow and as rodent models of drug use encompass greater depth and breadth, the integration of cross-species data with human GWAS may provide additional biological insight and help refine promising signals for functional follow-up.

## Acknowledgements

We would like to acknowledge the following funding sources that supported this work: NIDA DP1DA042103 (SBH and RHP), F32AA027435 (ECJ), Tobacco-Related Disease Research Program (TRDRP) Grant: #28IR-0070 (AAP), NIH Grants: P50DA037844 (AAP), AA018776 (ECJ) and P50 DA039841 (ECJ), K02DA032573 (AA), MH109532 (ECJ, AA and HJE).

## Authors contribution

SBH and ECJ performed analyses, made figures and tables and were responsible for writing the initial version of the manuscript. All other authors were instrumental in editing the manuscript and providing analytical advice for our study.

## REFERENCES

1. Goldman, D., Oroszi, G. & Ducci, F. The genetics of addictions: uncovering the genes. Nat. Rev. Genet. 6, 521–532 (2005).

2. Xian, H. et al. The heritability of failed smoking cessation and nicotine withdrawal in twins who smoked and attempted to quit. Nicotine Tob. Res. Off. J. Soc. Res. Nicotine Tob. 5, 245–254 (2003).

3. Wendt, F. R. et al. Natural selection influenced the genetic architecture of brain structure, behavioral and neuropsychiatric traits. bioRxiv 2020.02.26.966531 (2020). doi:10.1101/2020.02.26.966531

4. Quach, B. C. et al. Expanding the genetic architecture of nicotine dependence and its shared genetics with multiple traits. Nat. Commun. 11, 5562 (2020).

5. Liu, M. et al. Association studies of up to 1.2 million individuals yield new insights into the genetic etiology of tobacco and alcohol use. Nat. Genet. 51, 237–244 (2019).

6. Johnson, E. C. et al. A large-scale genome-wide association study meta-analysis of cannabis use disorder. The Lancet Psychiatry (2020). doi:10.1016/S2215-0366(20)30339-4

7. Zhou, H. et al. Association of OPRM1 Functional Coding Variant With Opioid Use Disorder: A Genome-Wide Association Study. JAMA Psychiatry 77, 1072–1080 (2020).

8. Kanehisa, M. & Goto, S. KEGG: kyoto encyclopedia of genes and genomes. Nucleic Acids Res. 28, 27–30 (2000).

9. Kanehisa, M., Furumichi, M., Tanabe, M., Sato, Y. & Morishima, K. KEGG: new perspectives on genomes, pathways, diseases and drugs. Nucleic Acids Res. 45, D353–D361 (2017).

10. González-Castro, T. B. et al. Identification of gene ontology and pathways implicated in suicide behavior: Systematic review and enrichment analysis of GWAS studies. Am. J. Med. Genet. Part B, Neuropsychiatr. Genet. Off. Publ. Int. Soc. Psychiatr. Genet. 180, 320–329 (2019).

11. Ferguson, L. B. et al. A Pathway-Based Genomic Approach to Identify Medications: Application to Alcohol Use Disorder. Brain Sci. 9, (2019).

12. Khatri, P., Sirota, M. & Butte, A. J. Ten Years of Pathway Analysis: Current Approaches and Outstanding Challenges. PLOS Comput. Biol. 8, e1002375 (2012).

13. Reynolds, T. et al. Interpretation of psychiatric genome-wide association studies with multispecies heterogeneous functional genomic data integration. Neuropsychopharmacology 46, 86–97 (2021).

14. Evans, L. M. et al. The Role of A Priori–Identified Addiction and Smoking Gene Sets in Smoking Behaviors. Nicotine Tob. Res. (2020).

15. Samson, H. H. & Czachowski, C. L. Behavioral measures of alcohol self-administration and intake control: rodent models. (2003).

16. Hopf, F. W. & Lesscher, H. M. B. Rodent models for compulsive alcohol intake. Alcohol 48, 253–264 (2014).

17. Spear, L. P. Consequences of adolescent use of alcohol and other drugs: studies using rodent models. Neurosci. Biobehav. Rev. 70, 228–243 (2016).

18. O’Dell, L. E. & Khroyan, T. V. Rodent models of nicotine reward: what do they tell us about tobacco abuse in humans? Pharmacol. Biochem. Behav. 91, 481–488 (2009).

19. Adkins, A. E. et al. Genomewide Association Study of Alcohol Dependence Identifies Risk Loci Altering Ethanol-Response Behaviors in Model Organisms. Alcohol. Clin. Exp. Res. 41, 911–928 (2017).

20. Palmer, R. H. C. et al. Multi-omic and multi-species meta-analyses of nicotine consumption. Transl. Psychiatry 11, 98 (2021).

21. Baker, E. J., Jay, J. J., Bubier, J. A., Langston, M. A. & Chesler, E. J. GeneWeaver: a web-based system for integrative functional genomics. Nucleic Acids Res. 40, D1067–D1076 (2011).

22. Durinck, S. et al. BioMart and Bioconductor: a powerful link between biological databases and microarray data analysis. Bioinformatics 21, 3439–3440 (2005).

23. Rodd, Z. A. et al. Differential gene expression in the nucleus accumbens with ethanol self-administration in inbred alcohol-preferring rats. Pharmacol. Biochem. Behav. 89, 481–498 (2008).

24. Philip, V. M. et al. High-throughput behavioral phenotyping in the expanded panel of BXD recombinant inbred strains. Genes. Brain. Behav. 9, 129–159 (2010).

25. Zhou, H. et al. Genome-wide meta-analysis of problematic alcohol use in 435,563 individuals yields insights into biology and relationships with other traits. Nat. Neurosci. (2020). doi:10.1038/s41593-020-0643-5

26. Walters, R. K. et al. Transancestral GWAS of alcohol dependence reveals common genetic underpinnings with psychiatric disorders. Nat. Neurosci. 21, 1656–1669 (2018).

27. Kranzler, H. R. et al. Genome-wide association study of alcohol consumption and use disorder in 274,424 individuals from multiple populations. Nat. Commun. 10, 1499 (2019).

28. Sanchez-Roige, S. et al. Genome-Wide Association Study Meta-Analysis of the Alcohol Use Disorders Identification Test (AUDIT) in Two Population-Based Cohorts. Am. J. Psychiatry (2018). doi:https://doi.org/10.1176/appi.ajp.2018.18040369

29. Finucane, H. K. et al. Partitioning heritability by functional annotation using genome-wide association summary statistics. Nat. Genet. 47, 1228–1235 (2015).

30. Bulik-Sullivan, B. K. et al. LD Score regression distinguishes confounding from polygenicity in genome-wide association studies. Nat. Genet. 47, 291–295 (2015).

31. Gandal, M. J., Leppa, V., Won, H., Parikshak, N. N. & Geschwind, D. H. The road to precision psychiatry: translating genetics into disease mechanisms. Nat. Neurosci. 19, 1397–1407 (2016).

32. Benjamini, Y. & Hochberg, Y. Controlling the false discovery rate: a practical and powerful approach to multiple testing. J. R. Stat. Soc. Ser. B 57, 289–300 (1995).

33. Zhou, H. et al. Meta-analysis of problematic alcohol use in 435,563 individuals identifies 29 risk variants and yields insights into biology, pleiotropy and causality. bioRxiv 738088 (2019). doi:10.1101/738088

34. Rhodes, J. S., Best, K., Belknap, J. K., Finn, D. A. & Crabbe, J. C. Evaluation of a simple model of ethanol drinking to intoxication in C57BL/6J mice. Physiol. Behav. 84, 53–63 (2005).

