## Supplementary_Information for "Genes identified in rodent studies of alcohol intake are enriched for heritability of human substance use"

*Non-substance use GWAS Samples*

To examine the specificity of the model organism alcohol and nicotine gene-sets, we also tested the enrichment of these gene-sets in several GWAS of non-substance use traits.

First, we selected GWAS summary statistics from previously published studies on both psychiatric and neurological disorders:

- **Schizophrenia:** N = 79,845^1^
  - Schizophrenia Working Group of the Psychiatric Genomics Consortium (2014). Biological insights from 108 schizophrenia-associated genetic loci. *Nature* **511**, 421 (2014).
- **Alzheimer’s disease:** N = 455,258^2^
  - Jansen, I. E. *et al.*(2019).Genome-wide meta-analysis identifies new loci and functional pathways influencing Alzheimer’s disease risk. *Nat. Genet.* **51**, 404–413 (2019).
- **Attention Deficit Hyperactivity Disorder (ADHD):** N = 55,374^3^
  - Demontis, D. *et al.* (2019). Discovery of the first genome-wide significant risk loci for attention deficit/hyperactivity disorder. *Nature genetics*, *51*(1), 63-75.
- **Autism Spectrum Disorder (ASD):** N = 173,773^4^
  - The Autism Spectrum Disorders Working Group of The Psychiatric Genomics Consortium (2017). Meta-analysis of GWAS of over 16,000 individuals with autism spectrum disorder highlights a novel locus at 10q24.32 and a significant overlap with schizophrenia." *Molecular autism* 8, 1-17.
- **Major Depressive Disorder (MDD):** N = 480,359^5^
  - Wray, N. R. *et al.* (2018). Genome-wide association analyses identify 44 risk variants and refine the genetic architecture of major depression. *Nature genetics*, *50*(5), 668-681.

Second, we selected GWASs that represented a range of non-substance use-related phenotypes. The GWAS were conducted on unrelated individuals of European descent and summary statistics were downloaded from the Neale lab UK Biobank GWASs (<http://www.nealelab.is/uk-biobank>). Traits were chosen if they had N > 50,000, significant heritability estimates, and no known connection to addiction traits:

- **Usual side of head for mobile phone use: right:** N = 303,009
- **Wears glasses or contact lenses:** N = 360,677
- **No lifetime tinnitus:** N = 117,882
- **Ankle spacing width:** N = 206,589
- **Hip circumference:** N = 360,521
- **Standing height:** N = 360,388
- **Wears glasses or contracs:** N = 360,677
- **Height (Standing):** N = 360,388
- **Bone Mineral Density (Heel):** N = 206,496
- **Diabetes (Diagnosed by Doctor):** N = 360,192
- **Ischemic Heart Disease (wide definition):** N = 361,194

**Supplementary Figure S1** Minimal Overlap Between Mouse and Rat Alcohol Gene-sets

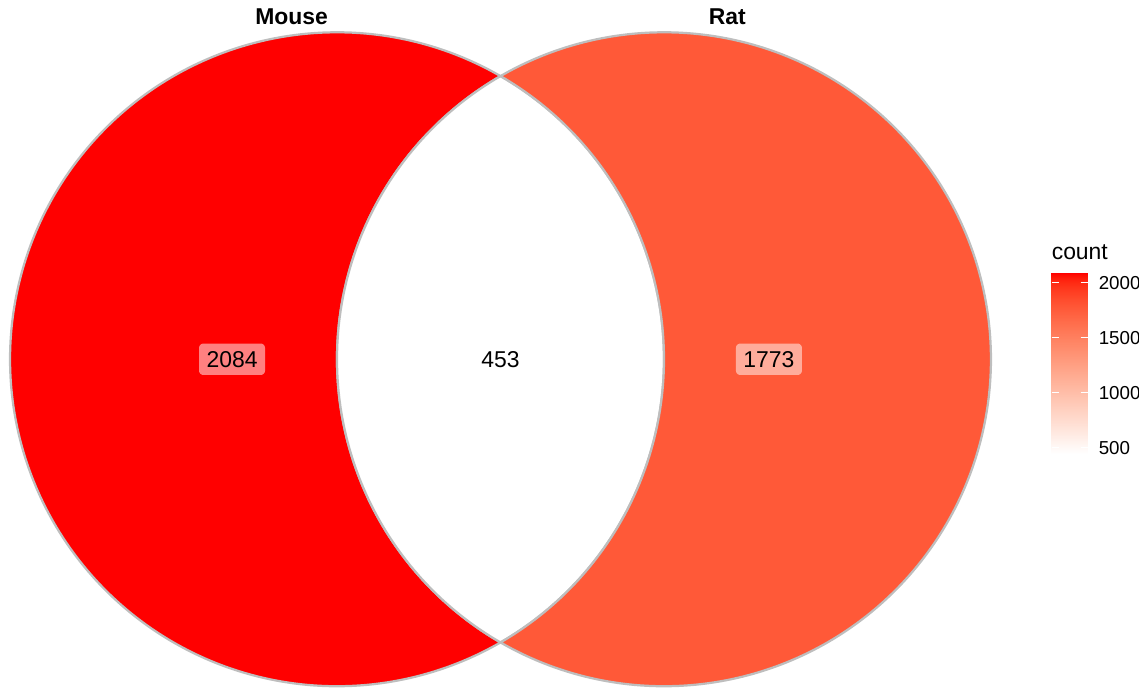

Gene-sets came from GeneWeaver.org, selecting Tier 3 brain-related gene expression studies from rodent paradigms of alcohol intake. In total rat and mouse gene-sets were comprised of 17 individual studies. For more information on these data see Supplementary Tables S1-S2.

**Supplementary Figure S2** Gene-set overlap in Mouse Paradigms of Alcohol Use

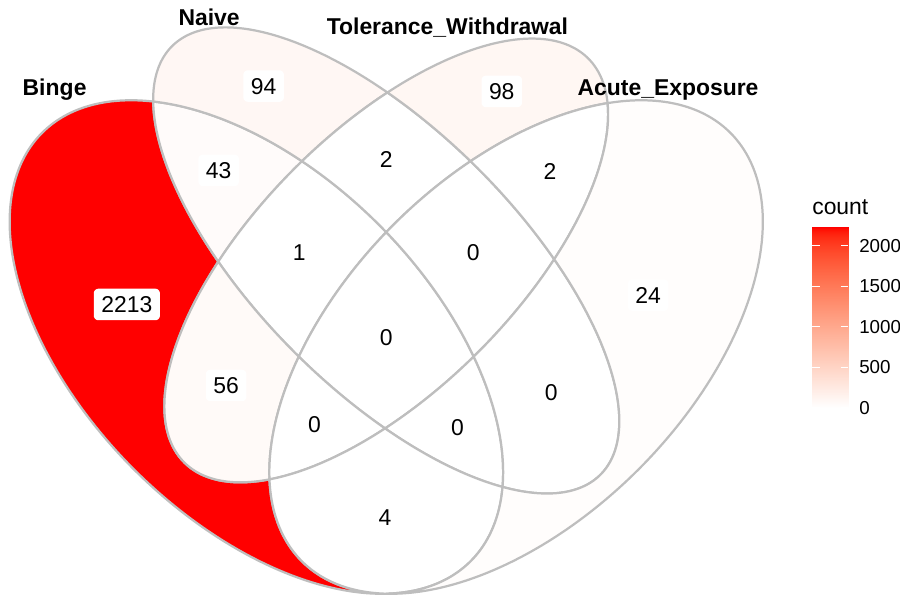

Naïve represents a paradigm where certain strains of mice consume or prefer alcohol more than a different genetic strain of mice. None of these mice were exposed to alcohol. Tolerance_withdrawal combines two paradigms of 1) acute functional tolerance and 2) withdrawal following acute and chronic alcohol use. Acute_Exposure includes a study that evaluated gene expression differences following a single (non-voluntary) treatment ethanol. Binge includes modified versions of the drinking in the dark paradigm where mice drink to intoxicating doses of ethanol. For more information on these genes see Supplementary Tables S1-S2.

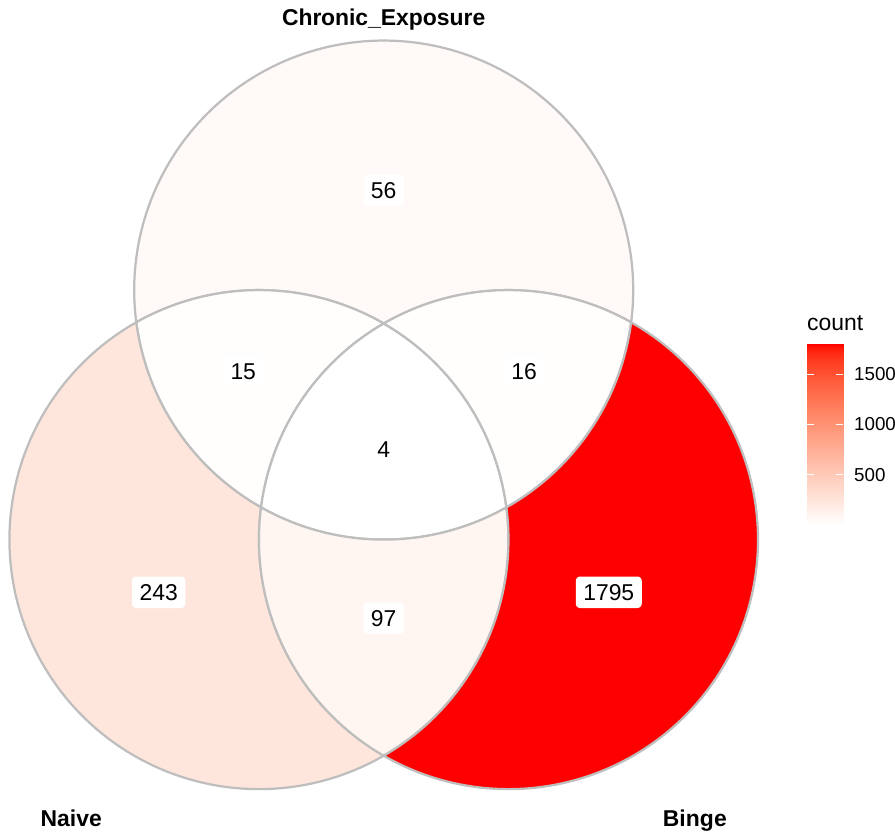
**Supplementary Figure S3** Gene-set overlap in Rat Paradigms of Alcohol Use

Naïve represents a paradigm where certain strains of rats consume or prefer alcohol more than a different genetic strain of rats. None of these rats were exposed to alcohol. Chronic_Exposure includes a study that evaluated gene expression differences following a multiple (non-voluntary) treatments of ethanol. Binge includes regular and inbred alcohol preferring rats that exhibit binge-like alcohol drinking. For more information on these genes see Supplementary Tables S1-S2.

**Supplementary Figure S4** Heritability enrichment of LD score regression’s conserved mammalian gene-set by GWAS trait

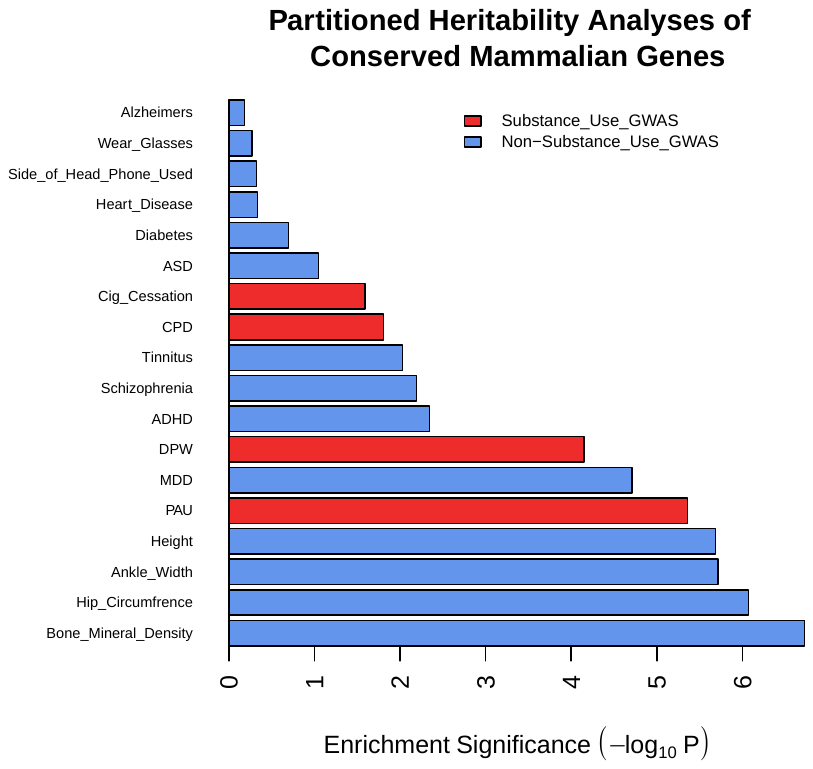

Barplot showing the significance of partitioned heritability analyses of the LD Score Regression conserved mammalian gene-set annotation. DPW = drinks per week, PAU = problematic alcohol use, CPD = cigarettes per day and Cig_Cessation = smoking cessation, ADHD = attention deficit hyperactivity disorder; MDD = major depressive disorder; ASD = autism spectrum disorder.

**Supplementary Figure S5** Significance of enrichment estimates increase with heritability among rodent substance use gene-sets.

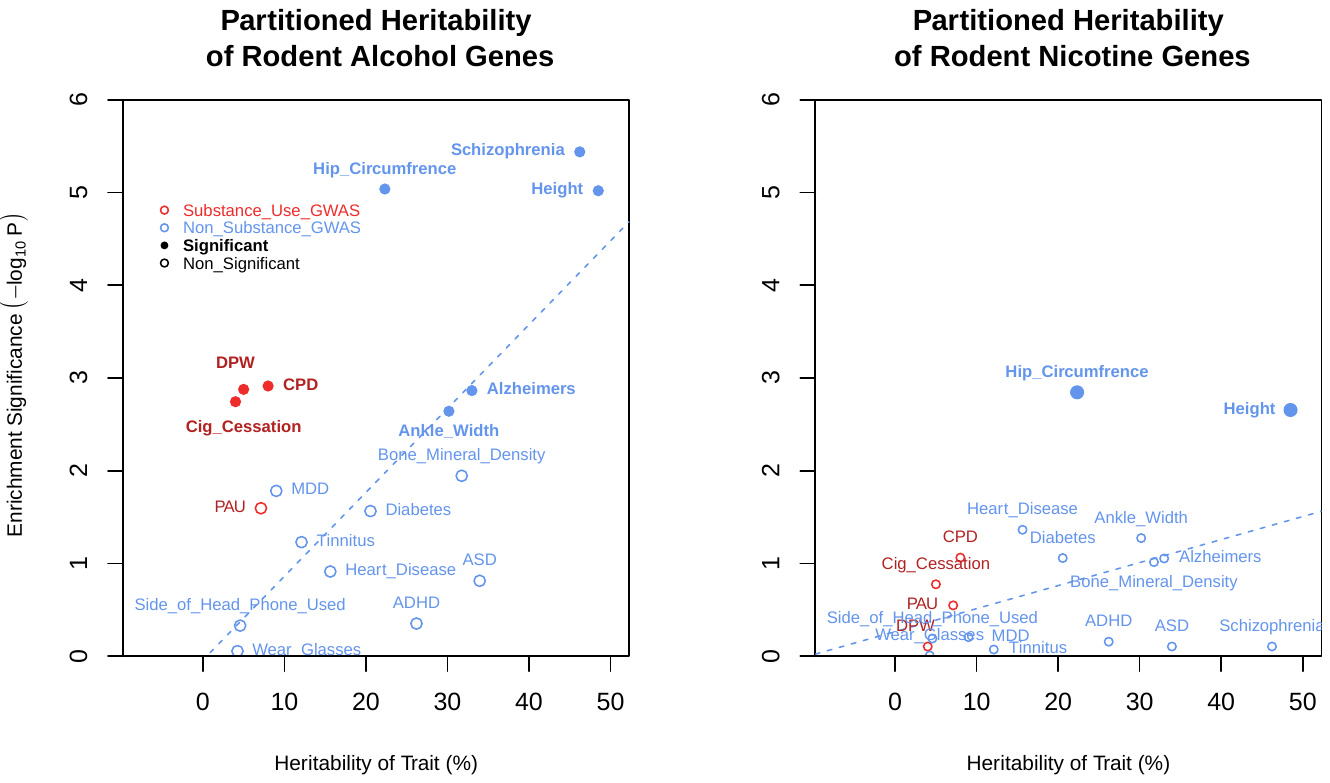

Overall, substance use traits tend to show more enrichment when compared with traits of similar heritability. The dashed blue line shows the best fitting regression line of a non-substance use trait’s heritability predicting the enrichment (Odds Ratio; OR) from partitioned heritability analyses. DPW = drinks per week, PAU = problematic alcohol use, CPD = cigarettes per day and Cig_Cessation = smoking cessation, ADHD = attention deficit hyperactivity disorder; MDD = major depressive disorder; ASD = autism spectrum disorder.

**Supplementary Figure S6** Correlations between partitioned heritability *significance* and trait heritabilities using rodent sucrose consumption and rodent locomotor behavior gene-sets

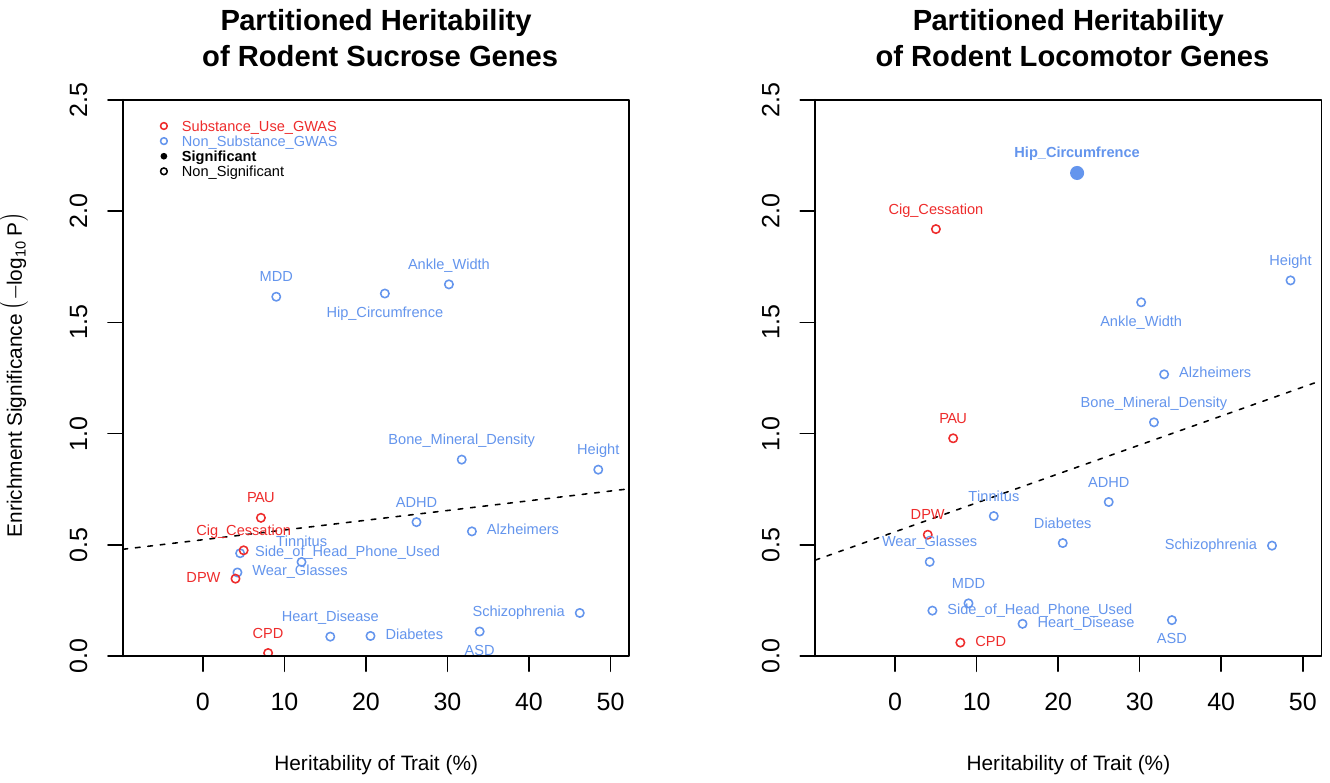

Plots show linear relationships between a trait's heritability and the *significance* (-log10 P-values) from partitioned heritability analyses using human GWASs adn rodent sucrose and locomotor gene-sets. Black dashed line shows the best fitting regression line among all traits. DPW = drinks per week, PAU = problematic alcohol use, CPD = cigarettes per day and Cig_Cessation = smoking cessation, ADHD = attention deficit hyperactivity disorder; MDD = major depressive disorder; ASD = autism spectrum disorder.

**Supplementary Figure S7** Associations between partitioned heritability *effect sizes* and trait heritabilities using rodent sucrose consumption and rodent locomotor behavior gene-sets

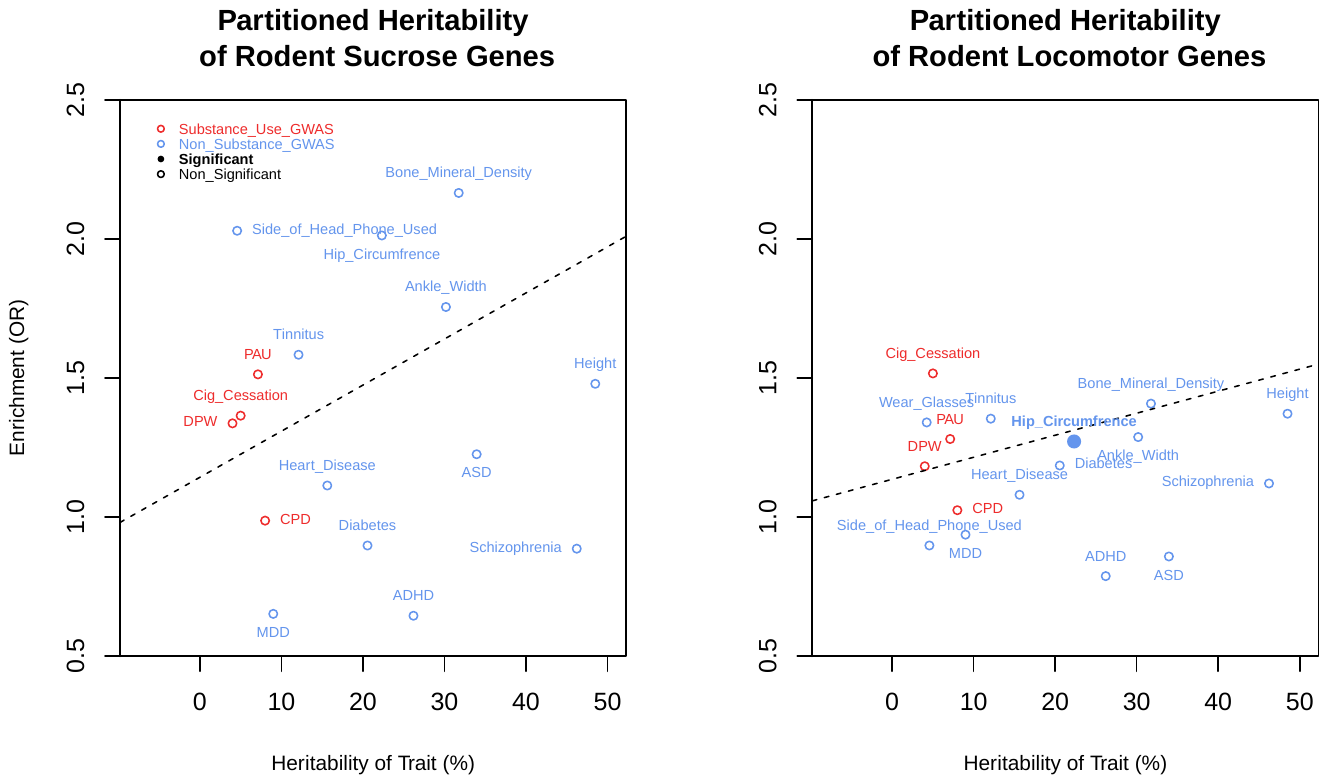

Plots show linear relationships between a trait's heritability and the *extent of enrichment* (Odds Ratio) from partitioned heritability analyses using human GWASs and rodent sucrose and locomotor gene-sets'. Black dashed line shows the best fitting regression line among all traits. DPW = drinks per week, PAU = problematic alcohol use, CPD = cigarettes per day and Cig_Cessation = smoking cessation, ADHD = attention deficit hyperactivity disorder; MDD = major depressive disorder; ASD = autism spectrum disorder.

**REFERENCES**

1.	Consortium, S. W. G. of the P. G. Biological insights from 108 schizophrenia-associated genetic loci. *Nature* **511**, 421 (2014).

2.	Jansen, I. E. *et al.* Genome-wide meta-analysis identifies new loci and functional pathways influencing Alzheimer’s disease risk. *Nat. Genet.* **51**, 404–413 (2019).

3.	Demontis, D., Walters, R. K., Martin, J., Mattheisen, M., Als, T. D., Agerbo, E., ... & Neale, B. M. (2019). Discovery of the first genome-wide significant risk loci for attention deficit/hyperactivity disorder. *Nature genetics*, *51*(1), 63-75.

4.	Meta-analysis of GWAS of over 16,000 individuals with autism spectrum disorder highlights a novel locus at 10q24. 32 and a significant overlap with schizophrenia." *Molecular autism* 8 (2017): 1-17

5.	Wray, N. R., Ripke, S., Mattheisen, M., Trzaskowski, M., Byrne, E. M., Abdellaoui, A., ... & Viktorin, A. (2018). Genome-wide association analyses identify 44 risk variants and refine the genetic architecture of major depression. *Nature genetics*, *50*(5), 668-681.
